# Suramin action in African trypanosomes involves a RuvB-like DNA helicase

**DOI:** 10.1101/2022.10.17.512644

**Authors:** Anna Albisetti, Silvan Hälg, Martin Zoltner, Pascal Mäser, Natalie Wiedemar

**Affiliations:** Swiss Tropical and Public Health Institute, 4051 Basel, Switzerland; University of Basel, 4002 Basel, Switzerland; Charles University in Prague, BIOCEV 252 50, Vestec, Czech Republic

**Author notes:** Equal contribution (Pascal Mäser, Natalie Wiedemar). Corresponding author: Address Swiss Tropical and Public Health Institute, Kreuzstrasse 2, 4123 Allschwil Switzerland, Mail. Present address: University of Dundee, Wellcome Centre for Anti-Infectives Research, Dundee DD1 5EH (Natalie Wiedemar).

## Abstract

Suramin is one of the oldest drugs in use today. It is still the treatment of choice for the hemolymphatic stage of African sleeping sickness caused by *Trypanosoma brucei rhodesiense* and it is also used for surra in camels, caused by *Trypanosoma evansi*. Yet despite one hundred years of use, suramin’s mode of action is not fully understood. Suramin is a polypharmacologic molecule that inhibits diverse proteins. Here we demonstrate that a DNA helicase of the pontin/ruvB-like 1 family, termed *T. brucei* RuvBL1, is involved in suramin resistance in African trypanosomes. Bloodstream-form *T. b. rhodesiense* under long-term selection for suramin resistance acquired a homozygous point mutation, isoleucin-312 to valine, close to the ATP binding site of *T. brucei* RuvBL1. The introduction of this missense mutation, by reverse genetics, into drug-sensitive trypanosomes significantly decreased their sensitivity to suramin. Intriguingly, the corresponding residue of *T. evansi* RuvBL1 was found mutated in a suramin-resistant field isolate, in that case to a leucin. RuvBL1 (Tb927.4.1270) is predicted to build a heterohexameric complex with RuvBL2 (Tb927.4.2000). RNAi-mediated silencing of gene expression of either *T. brucei* RuvBL1 or RuvBL2 caused cell death within 72 h. At 36 h after induction of RNAi, bloodstream-form trypanosomes exhibited a cytokinesis defect resulting in the accumulation of cells with two nuclei and two or more kinetoplasts. Taken together, these data indicate that RuvBL1 DNA helicase is among the primary targets of suramin in African trypanosomes.

**Abstract Importance:** African trypanosomes cause sleeping sickness in humans, nagana in cattle, and surra in camels – lethal diseases for which there is no vaccine and only few drugs. One of the drugs is suramin, developed by Bayer in 1916. Yet despite 100 years of use, suramin’s mode of action is not fully understood at the molecular level. Here we show that a DNA helicase is involved: *Trypanosoma brucei rhodesiense* (causative agent of sleeping sickness) selected for suramin resistance acquired a point mutation in the DNA helicase RuvBL1 that, when introduced to wild-type trypanosomes, reduced their sensitivity to suramin. Intriguingly, the same site in RuvBL1 was mutated also in a suramin-resistant field isolate of *T. evansi* (causative agent of surra). We further demonstrate that RuvBL1 is essential for proper cell division of *T. brucei*. Thus we conclude that inhibition of RuvBL1 contributes to the trypanocidal action of suramin.

## Introduction

Suramin is an enigmatic molecule. It has a molecular weight of 1300 Da, carries six negative charges at physiological pH, and it is not orally bioavailable – yet despite this lack of drug-like properties, suramin has been in use as an antiinfective agent for over a century, in human as well as in veterinary medicine. Its primary indications are human African trypanosomiasis (HAT) and surra in camels (Giordani, 2016); suramin was also used for the treatment of onchocerciasis, caused by the nematode *Onchocerca volvulus* (Hawking, 1978). Suramin is a polypharmacologic molecule that inhibits dozens of different enzymes and receptors (Voogd, 1993). Its potential applications in medicine are manifold (Wiedemar, 2020), ranging from antiparasitic, antiviral, and anticancer chemotherapy to the use as an antidote for snakebite (Murakami, 2005) or as a new treatment option for autism (Naviaux, 2017). Paradoxically, we know more about the mechanisms of action of suramin for these repurposed applications than for its primary indication, African trypanosomiasis.

*Trypanosoma brucei* subspecies and *Trypanosoma evansi* cause neglected tropical diseases affecting humans and livestock. Human African trypanosomiasis is largely under control thanks to successful programs against sleeping sickness and tsetse flies, and new drugs fed into the development pipeline, in particular fexinidazole as a new, oral treatment for *T. b. gambiense* infections (Dickie, 2020; Franco, 2020). The animal trypanosomiases, in contrast, remain a serious problem for agriculture, causing huge economic losses and hindering development (Giordani, 2016). As *T. evansi* is not restricted to tsetse flies for transmission, it has a wider geographical distribution than *T. brucei*. This makes it a problem not only for Africa but also for countries in Asia, South America and potentially Europe (Desquesnes, 2013). Suramin is still being used to treat first-stage (i.e. hemolymphatic) *T. b. rhodesiense* infections in humans, and it was used extensively for *T. evansi* infections in livestock until suramin resistance became widespread (El Rayah, 1999; Zhou, 2004).

The molecular nature of the suramin target(s) in African trypanosomes has been the subject of many studies. Suramin impairs oxygen consumption and ATP production in *T. brucei* bloodstream forms (Fairlamb, 1980), and it inhibits several of the glycolytic enzymes of *T. brucei*: hexokinase, aldolase, glycerol-3-phosphate dehydrogenase, and phosphoglycerate kinase (Willson, 1993). Suramin was also shown to inhibit other trypanosomal enzymes such as glycerophosphate oxidase (Fairlamb, 1977), serine oligopeptidase (Morty, 1998), and RNA editing ligase (Zimmermann, 2016). If all these enzymes are susceptible to suramin, which is the primary target? It is still currently unclear which of the *T. brucei* enzymes that are inhibited by suramin are responsible for the cytotoxic effect of the drug. These are expected (i) to be essential for the proliferation of bloodstream-form trypanosomes and (ii) to be affected in suramin-resistant mutants.

We have previously attempted to elucidate the mechanism of action of suramin through the generation of resistant *T. brucei* mutants. *In vitro* selection at high concentrations of suramin (25-fold the IC_50_) readily produced derivatives with a 100-fold lower susceptibility to suramin (Wiedemar, 2018). However, this approach provided insights into the uptake mechanisms of suramin rather than its targets (Wiedemar, 2019). The suramin-resistant trypanosomes expressed a specific variant surface glycoprotein (VSG) variant termed VSG^Sur^, which binds suramin with high affinity (Zeelen, 2021) and provides an alternative endocytosis route to the known endocytosis pathways via either ISG75 (Alsford, 2012) or low-density lipoprotein (Vansterkenburg, 1993). Here we have further selected VSG^Sur^-expressing *T. brucei* for high-level suramin resistance, which has led to the identification of a DNA helicase as a new, and likely primary, target of suramin in African trypanosomes.

## Results

### Selection of a 1000-fold suramin resistant T. b. rhodesiense mutant

Starting from a fresh clone of our drug testing reference strain, *T. b. rhodesiense* STIB900_c1, we had previously generated several suramin-resistant lines that are resistant because they express VSG^Sur^, a peculiar variant surface glycoprotein that binds suramin (Wiedemar, 2018; Wiedemar, 2019; Zeelen, 2021). One of these lines, *T. b. rhodesiense* STIB900_c1_sur1, was subjected to further selection. Bloodstream-form cultures were continuously exposed to sublethal concentrations of suramin over a period of one year. Starting from 1 μM, the concentration of suramin was gradually increased to 18.9 μM as the trypanosomes were losing their susceptibility (Figure 1A). The finally obtained derivative, *T. b. rhodesiense* STIB900_c1_sur1_18900, had a 50% inhibitory concentration (IC_50_) to suramin of 11.0 μM, more than a thousand-fold higher than that of the parental clone *T. b. rhodesiense* STIB900_c1 (Fig. 1B). With progressing suramin resistance the trypanosomes also lost their susceptibility to the related molecule trypan blue, albeit to a lesser extent than observed for suramin (Fig. 1B). No alteration was observed regarding the sensitivity to the reference drugs melarsoprol and pentamidine (Fig. 1B).

**Figure 1.**
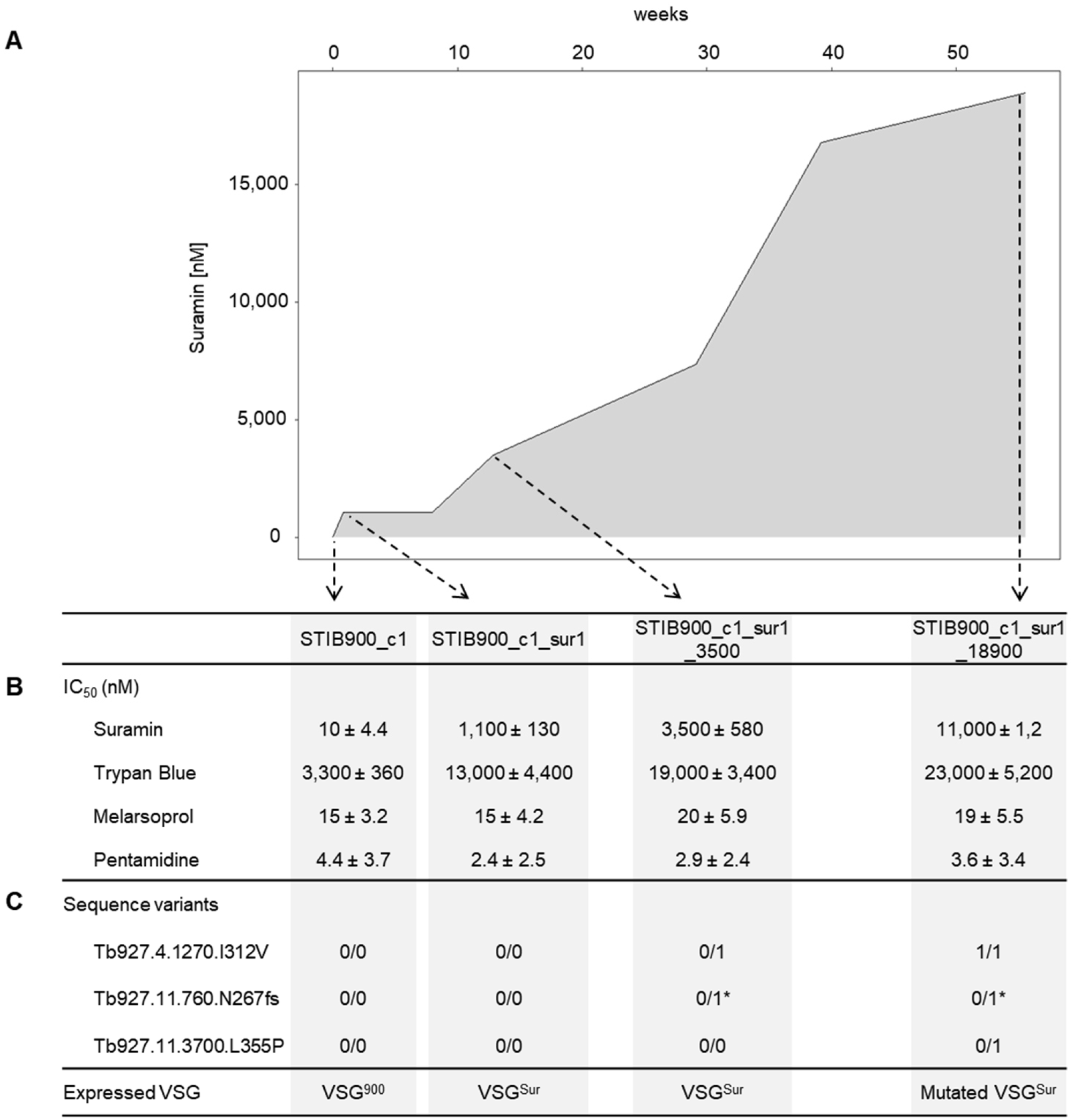
Suramin-selected *T. b. rhodesiense* STIB900 derivatives, their phenotypes and genotypes. (A) Time course and suramin concentrations used for the selection of STIB900-c1. (B) Drug sensitivity of the STIB900-c1 derivatives at different time points during suramin selection. (C) Genes that acquired non-synonymous mutations during the selection (0/1, heterozygous mutation; 1/1, homozygous mutation; asterisk, only 30% of reads mutated). VSG-Sur and the mutated version of VSG-Sur have been described previously (Wiedemar et al., 2018; Zeelen et al., 2021).

### Candidate genes for suramin resistance and mutations therein

Given the role that VSG can play in suramin resistance (Wiedemar, 2018; Wiedemar, 2019; Zeelen, 2021), we first determined which VSG was expressed by the suramin-resistant lines. For this purpose, the VSG coding regions were amplified by PCR from reverse-transcribed mRNA. As reported previously (Wiedemar, 2018), *T. b. rhodesiense* STIB900_c1_sur1 had switched from VSG^900^ to VSG^Sur^. The same VSG^Sur^ was expressed also by the intermediate line *T. b. rhodesiense*_c1_sur1_3500 (Fig. 1C). The final line *T. b. rhodesiense*_c1_sur1_18900, however, had acquired 14 point mutations in the expressed VSG^Sur^, 8 of which were non-synonymous. These mutations and their enhancing effect on suramin resistance were reported elsewhere (Zeelen, 2021).

Aiming to identify further genetic mutations associated with suramin resistance, in particular mutations of genes other than VSG, we performed RNA-Seq of the sensitive parental clone (*T. b. rhodesiense* STIB900_c1), its VSG^Sur^-expressing derivative (STIB900_c1_sur1), and the two derivatives with higher-level suramin resistance (STIB900_c1_sur1_3500 and STIB900_c1_sur1_18900). All cultures were grown in the absence of suramin. Mapping of the obtained reads revealed three non-synonymous mutations in non-*VSG* genes that had emerged during the selection process. None of these variants are annotated as known nucleotide polymorphisms in the TriTryp SNP database (tritrypdb.org) (Aslett, 2010).

Two of the polymorphisms were heterozygous (Fig. 1C). The first was an insertion at nucleotide 798 in the gene for a putative phosphatase 2C (Tb927.11.760), leading to a frameshift from amino acid 267 onwards. However, fewer than 30% of the RNA-Seq reads carried this mutation, so it was not further investigated. The second was a missense mutation (t1064c) in the gene for an ADAM Cysteine-Rich Domain containing protein of the YagE family (Tb927.11.3700), resulting in the mutation leucine to proline at position 355. However, the mutation was present only in *T. b. rhodesiense* STIB900_c1_sur1_18900 (Fig. 1C), and the *T. brucei* YagE orthologue is not essential in bloodstream-form trypanosomes as determined by the genome-wide RNAi screen (Alsford, 2011). Therefore, again, the gene was not further investigated here. The third mutation was the most interesting because it was heterozygous in the intermediate line *T. b. rhodesiense* STIB900_c1_sur1_3500 and homozygous in the final line STIB900_c1_sur1_18900 (Fig. 1C). This was confirmed by PCR on genomic DNA followed by Sanger sequencing. The mutation was a missense variant (a934g) in the RuvB-like 1 DNA helicase gene (Tb927.4.1270; herein called RuvBL1), resulting in the amino acid substitution of isoleucine-312 with valine (I312V) in the *T. brucei* RuvBL1 orthologue.

### In situ mutation of RuvBL1 in T. brucei causes suramin resistance

First, we investigated the effect of the RuvBL1 I312V mutation in the genetic background it originally appeared in. The parental line *T. b. rhodesiense* STIB900_c1_sur1, which is suramin resistant but homozygous for wild-type RuvBL1 (Fig. 1), was transfected with a construct encoding for the I312V mutant of RuvBL1 (Fig. 2A). This gave rise to only one positive clone that had the construct correctly integrated into the *RuvBL1* locus by homologous recombination. The transfected clone was heterozygous for the mutation as determined by PCR and Sanger sequencing, and it exhibited a 2.2-fold increase in suramin resistance as compared to untransfected *T. b. rhodesiense* STIB900_c1_sur1 (Fig 2B). The IC_50_ shift was slightly lower than the 2.9-fold increase observed for *T. b. rhodesiense* STIB900_c1_ sur1_3500 (Fig. 2B), the line in which the heterozygous mutation I312V in RuvBL1 had emerged during suramin selection (Fig. 1). Thus, there might be additional factors contributing to suramin resistance in the laboratory-evolved line.

**Figure 2.**
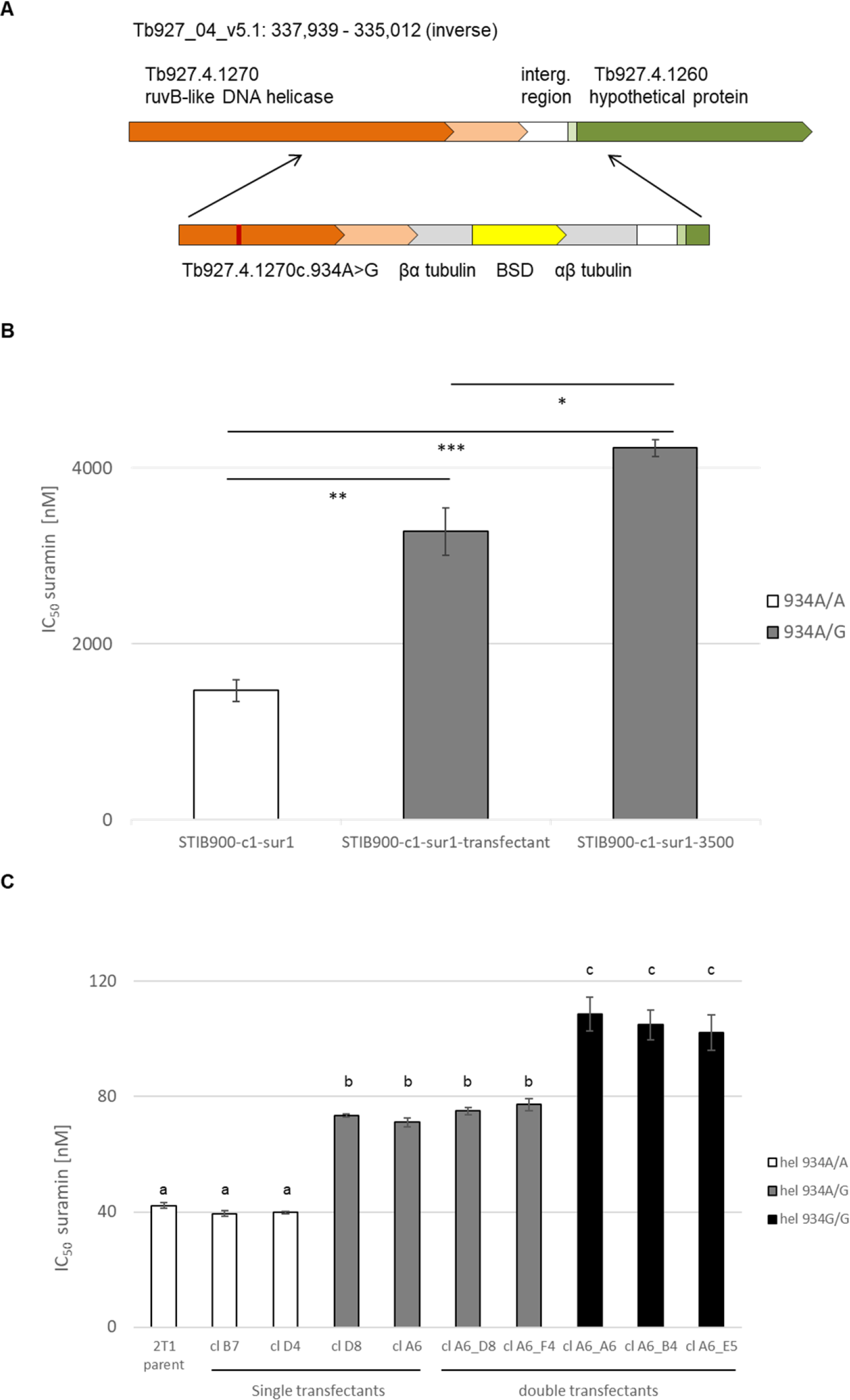
*In situ* introduction of mutated RuvB-like helicase 1 causes suramin resistance. (A) Genomic context and the construct for reverse genetics, which consists of the last 704 nucleotides of coding sequence and the 3’UTR of *RuvBL1*, a βα-tubulin mRNA processing site, a blasticidine resistance gene (BSD), an αβ-tubulin mRNA processing site; and 309 nt homologuous region downstream of the gene. The location of the mutation a934g, as part of the Val^312^ codon, is marked as dark red line. (B) Suramin sensitivities of parental *T. b. rhodesiense* STIB900-c1-sur1, a *RuvBL1-*Val^312^ transfectant thereof, and the suramin selected line STIB900-c1-sur1_3500 represented as 50% inhibitory concentrations (IC_50_). N = 3 biological replicates, each consisting of 2 technical replicates. Error bars represent standard error. P-values as calculated with Tukey’s multiple comparison test, * p<0.05; ** p<0.001; *** p<0.0001. (C) Suramin sensitivities of *T. b. brucei* 2T1 clones after single or double transfection with mutated *RuvBL1*. The clones after the first transfection were either heterozygous for the mutated helicase (D8 and A6; gray) or had remained wildtype (B7 and D4; white). After the second transfection, the clones were either homozygous (A6_A6, A6_B4 and A6_E5; black) or had remained heterozygous (A6_D8 and A6_F4; gray). Error bars represent standard error. N = 4 biological replicates, each consisting of 2 technical replicates, N = 8 for 2T1 and cl. A6. Significant differences between groups (p < 0.001) as calculated with One-Way Anova followed by Tukey’s multiple comparison test are indicated in lowercase.

To test the effect of the mutation I312V in a neutral genetic background, the same construct (Fig. 2A) was transfected into *T. b. brucei* 2T1 bloodstream forms (Alsford, 2005). This generated two positive clones that were heterozygous for the mutation. In addition, two clones were obtained which had integrated the selective marker (the blasticidin S deaminase gene) but not the mutation in RuvBL1, presumably due to a recombination event downstream of codon 312. These two clones were used as additional controls. The IC_50_ values for suramin in the heterozygous RuvBL1 I312V mutants were 1.8-fold higher than those of the controls (Fig 2C), while no significant alteration was observed regarding the susceptibility to trypan blue, melarsoprol, or pentamidine (Suppl. Fig. S1).

Homozygous *T. b. brucei* mutants were generated after exchanging the blasticidin resistance marker of the construct with the neomycin S transferase gene, transfecting one of the heterozygous transgenic clones, and selection for clones that were resistant to blasticidin as well as G418. Out of five positive clones, three were homozygous for valine-312 while two had remained heterozygous. The former exhibited a further 1.4-fold drop in suramin sensitivity as compared to the latter, and a 2.4-fold drop as compared to parental *T. b. brucei* 2T1 (Fig. 2C). Again, the sensitivity to trypan blue, melarsoprol, and pentamidine remained unaffected (Fig S1). Thus, the mutation I312V in the RuvBL1 ortholog of *T. brucei* causes a resistance phenotype that is specific for suramin. This strongly indicates that the *T. brucei* RuvBL1 helicase is involved in the mode of action of suramin.

### A T. evansi suramin-resistant field isolate carries a similar mutation in RuvBL1 helicase

Suramin resistance is a major problem in the control of surra in camels (El Rayah, 1999; Zhou, 2004). We had generated whole-genome sequencing data of a suramin-resistant *T. evansi* field isolate, Westry2 (El Rayah, 1999), which we inspected for the orthologues of the identified candidate genes. In particular, we looked for non-synonymous deviations from the reference sequence *T. evansi* STIB805 (Carnes, 2015) that were not present in a suramin-sensitive *T. evansi* isolate, STIB779 (El Rayah, 1999). The RuvBL1 sequences of the suramin-sensitive *T. evansi* STIB779 and STIB805 (TevSTIB805.4.1310) were identical to that of *T. brucei*. Intriguingly, however, *T. evansi* Westry2 carried a heterozygous mutation in the RuvBL1 gene at exactly the same position as the laboratory-selected *T. b. rhodesiense* line, i.e. at codon 312. The mutation was a934t, leading to an amino acid replacement from isoleucine to leucine (I312L).

### Isoleucin-312 is located close to the active site of RuvBL1

Tb927.4.1270 shares 65% sequence identity with *Saccharomyces cerevisiae* RuvB-like 1 DNA helicase. However, despite a high degree of sequence conservation among the eukaryotic RuvB-like 1 proteins, the orthologues from the kinetoplastids form a clearly distinct branch in the phylogenetic tree (Fig. 3A). RuvBL1 forms a heterohexameric complex with the paralogous RuvB-like DNA helicase RuvBL2. The corresponding homolog in *T. brucei*, Tb927.4.2000, is encoded 170 kb upstream of RuvBL1 on chromosome 4. The RuvBL complex facilitates ATP-dependent nucleosome sliding as part of larger complexes with a function in nucleosome remodeling in eukaryotes. In yeast the two distinct chromatin remodelling complexes, INO80 and SWR1, are well characterized (Gerhold, 2014). A SWR1-like remodeller complex in *T. brucei* was described recently (Vellmer, 2022). Homology models of RuvBL1 and RuvBL2 were built with high confidence using *S. cerevisiae* orthologs (pdb 6GEN) (Willhoft, 2018) as template. I312 in the RuvBL1 structural model (labeled in orange in Fig. 3B) is located downstream of the ATP-binding domain, six residues upstream the Walker B motif (Matias, 2006). The latter motif is highly conserved (Fig. 3C, Suppl. Fig. S2) and has, together with the Walker A motif, a key role in ATP binding and hydrolysis by coordinating the β and γ phosphates of ATP and the water activating magnesium ion (Wendler, 2012). Bound ATP and the I312 side chain are more than 23 Å apart, rendering it unlikely but not impossible, that suramin, whose structure spans approximately 30 Å, interacts directly with the ATP binding site. Superimposing the homology models to the *S. cerevisiae* RuvBL heterohexameric assembly (Fig. 3D) implies that the mutant position is close to the subunit interface and the central pore, a position where suramin binding could interfere with protein-protein or protein-DNA interaction. Lastly, I312 precedes a region neighbouring the nucleotide interacting sensor 1 motif (Fig. 3B), which has been implicated in substrate sensing (Erzberger, 2006).

**Figure 3.**
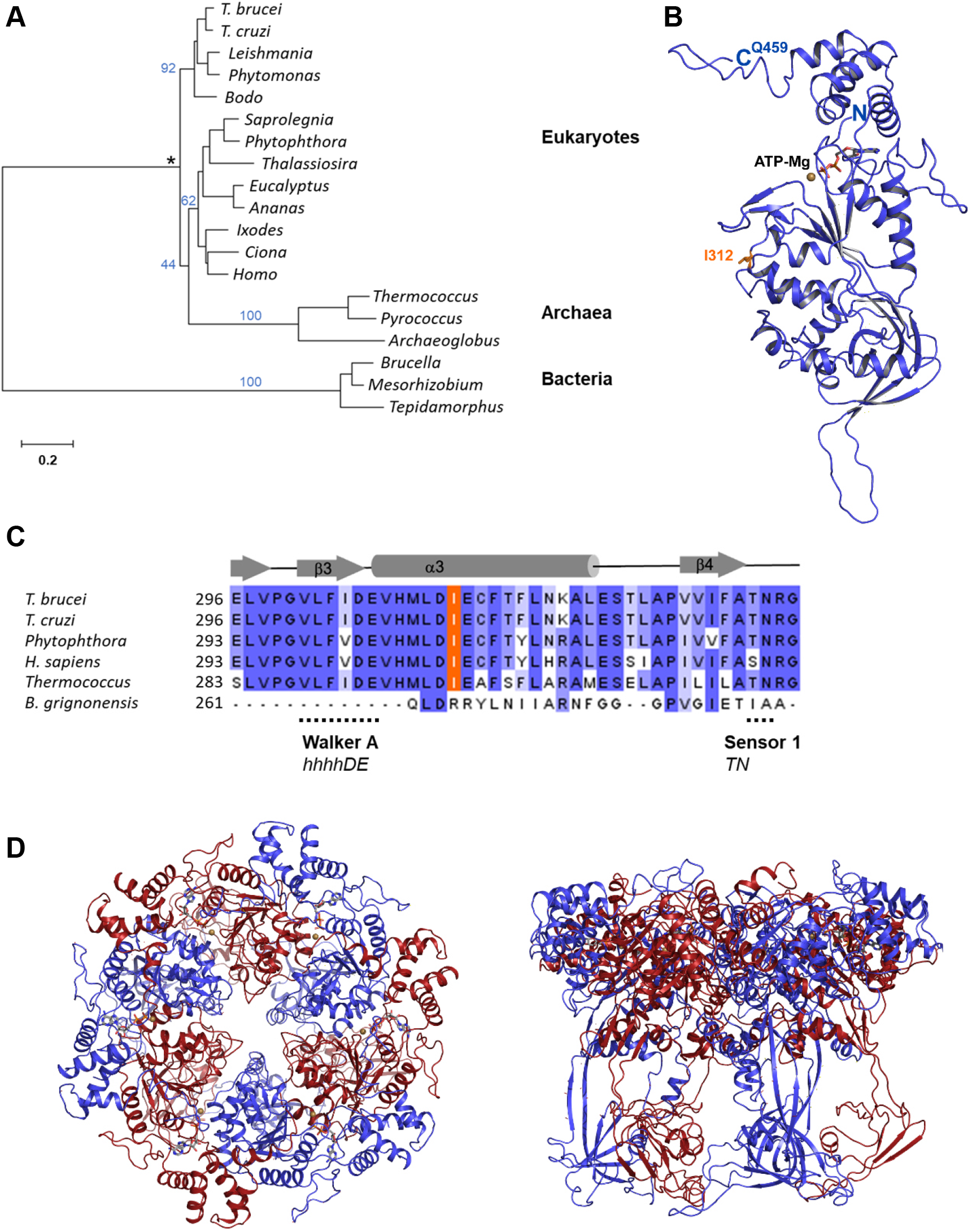
Phylogenetic analysis of *T. brucei* RuvB-like and homology model. (A) Neighbor-Joining phylogenetic tree of RuvB-like helicases from different taxonomic groups showing how the kinetoplastid sequences form a distinct clade (blue numbers, percent positives of 1000 rounds of bootstrapping; asterisk, presumed root based on the prokaryotic outgroup; scale bar, number of amino acid substitutions per site; for the complete sequence alignment see Fig. S2). (B) Homology model of TbRuvBL1 (Tb927.4.1270) in cartoon representation with Ile^312^ drawn in purple sticks. ATP (sticks) and Mg^2+^ (sans colored sphere) are superimposed from the *S. cerevisiae* RuvBL1 structure (pdb ID 6GEN) (Willhoft, 2018). (C) Section of a multiple sequence alignment of RuvB-like helicases showing how Ile^312^ is conserved across species (the full alignment is in Figure S2). The location of Walker B motif (hhhhDE) (Hanson, 2005; Matias, 2006) and the nucleotide interacting sensor 1 motif are indicated. (D) Homology model of the heterohexameric complex of TbRuvBL1 (Tb927.4.1270, blue) and TbRuvBL2 (Tb927.4.2000, red) in cartoon representation, top and side view.

### Overexpression of RuvB-like helicases only marginally affects suramin resistance

Inhibition of a target protein by a drug can, in certain cases, be compensated for by overexpression of the target itself. To test whether overproduction of RuvBL1 helicase leads to suramin resistance, we transfected *T. b. brucei* 2T1 cells with a pRPa-based construct for tetracycline-inducible overexpression of *RuvBL1*. Three positive transfectants were investigated. Induction with tetracycline (1 μg/mL) led to an increase in *RuvBL1* mRNA levels of 2.2 to 3.0 fold as measured by qPCR, but the suramin sensitivity in the induced cells was not altered as compared to uninduced cells (Suppl. Fig. S3A and S3B). Overproduction of RuvBL1 alone might not have an effect because the protein is predicted to build a heteromeric complex together with RuvBL2 (Fig. 3A). To simultaneously overexpress both genes, we made a construct containing *RuvBL1* and *RuvBL2* separated by an αβ tubulin mRNA processing site. Again, *T. b. brucei* 2T1 cells were transfected and three positive clones were obtained and investigated.

Overexpression of the target genes was confirmed in principle, but it happened to a much lower extent than for *RuvBL1* alone: after induction by tetracycline, the mRNA levels of *RuvBL1* and *RuvBL2* only rose 1.3- to 1.6-fold and 1.3- to 1.5-fold, respectively (Suppl. Fig. S3C). The tetracycline-induced cells had a slightly higher IC_50_ to suramin than the uninduced cells for all three clones, but the difference was not statistically significant (Suppl. Fig. S3D). So overall, the experiments linking overexpression of RuvBL orthologues in *T. brucei* to suramin-sensitivity were not conclusive. This is not surprising: the expression levels of the *RuvBL* orthologues were only slightly increased, and even if it were possible to reach high-level overexpression of *RubBL1* and *RubBL2*, this would unlikely result in the overproduction of a functional complex. Additional factors are necessary for function and proper folding to the predicted ring-like structure (Abrahao, 2021).

### The RuvBL1 and RuvBL2 helicases are both independently essential for T. brucei

If RuvBL1 is indeed a drug target, then it has to be essential for the proliferation of bloodstream-form trypanosomes. Using pRPa-iSL-based plasmids (Alsford, 2005), we generated two stem-loop RNAi cell lines for tetracycline-inducible silencing of *RuvBL1* or *RuvBL2* expression in bloodstream-form *T. brucei*. Both lines showed a strong down-regulation of *RuvBL1* and *RuvBL2* expression: 24 h after addition of tetracycline (1 μg/mL), the mRNA levels of either gene had dropped to below 20% of the original value as determined by qPCR (Fig 4A). Both individual genetic knock-downs led to a rapid slowdown of growth within the first 24 h, which culminated into growth arrest and cell death within three days after induction (Fig 4B). These observations confirmed the results of the high-throughput RNAi screen in *T. brucei* (Alsford, 2011), in which *RuvBL1* and *RuvBL2* were both categorized as essential genes for bloodstream-form parasites.

**Figure 4.**
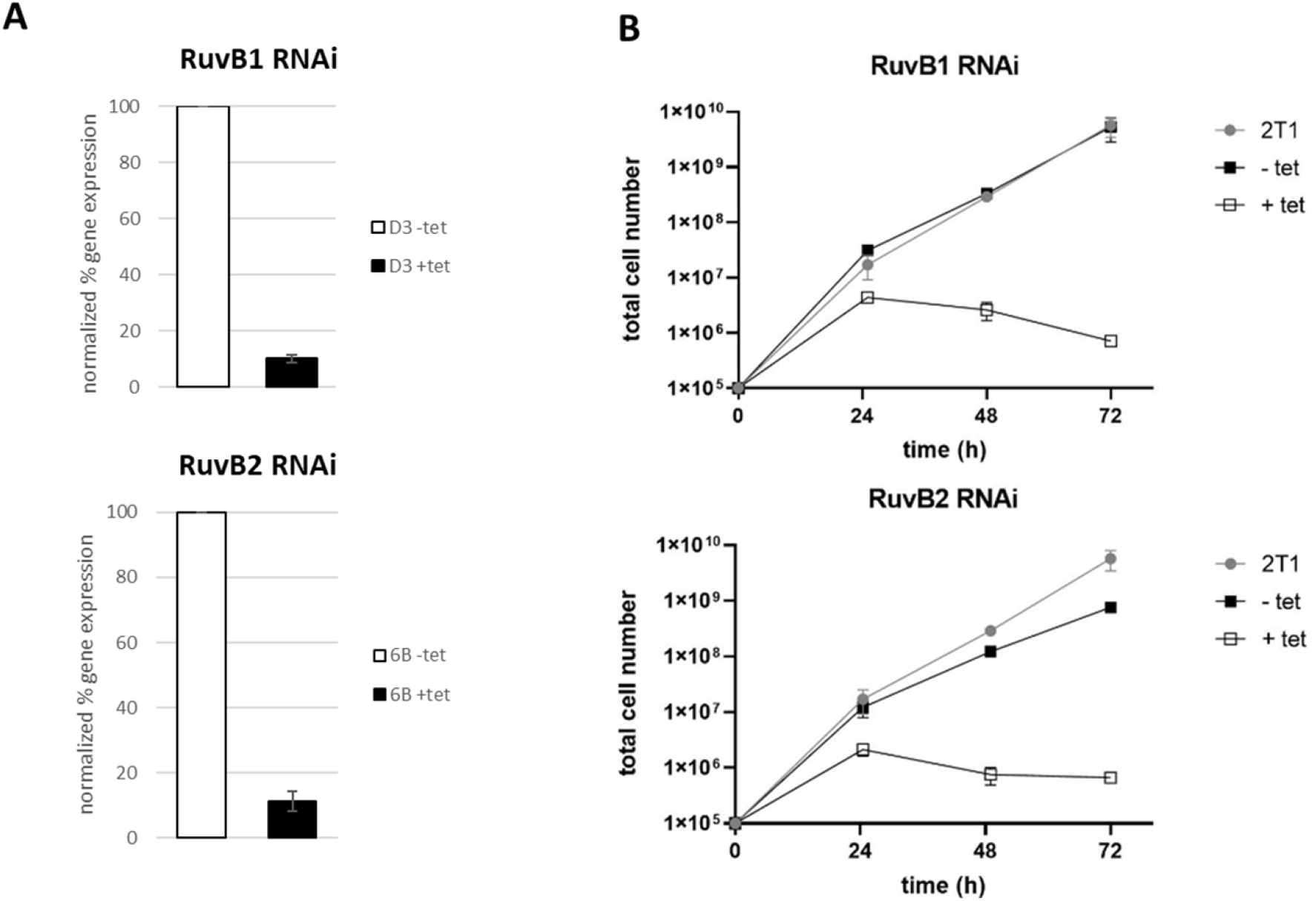
RuvBL1 and RuvBL2 are essential for bloodstream-form *T. brucei*. (A) Steady-state gene expression levels as determined with qPCR at 24 h post tetracycline (tet) induction (n=3, with 2 technical replicates). (B) Representative growth curves of uninduced (−tet) and induced clone (+tet) compared to the parental line *T. b. brucei* 2T1 (n=3). Upper panel, transfectant clone D3 – *RuvBL1* RNAi; lower panel, transfectant 6B – *RuvBL2* RNAi.

### Downregulation of either RuvBL1 or RuvBL2 blocks cytokinesis

Due to the rapid cell growth arrest upon down-regulation of either RuvB-like helicase, we investigated the DNA content of the RNAi lines in the short time window before the trypanosomes were dying. The cell lines were induced with 1 μg/mL tetracycline for 24 h and 36 h, fixed, and 4’,6-diamidino-2-phenylindole (DAPI)-stained to visualize the nuclei (N) and the kinetoplasts (K; the DNA-containing structure within the single mitochondrion). The parental line (*T. b. brucei* 2T1), the uninduced, and the induced RNAi cell lines were analyzed by light microscopy regarding their DNA content. Compared to the typical wildtype phenotypes in the parental line (mostly 1K1N cells with one kinetoplast and one nucleus, plus some cells with 2K1N and 2K2N; black bars in Fig. 5A), a marked decrease of the 1K1N and 2K1N phenotypes was observed in the tetracycline-induced lines (grey bars in Fig. 5A) in favor of a clear accumulation of 2K2N cells. At 36 h post-induction, the 1K1N cells had dropped from 80% of the total population to about 20% in the *RuvBL1* RNAi line, and from 76% to 30% in *RuvBL2* RNAi line. At the same time, the 2K2N cells had increased from 7% to 37% in the *RuvBL1* RNAi line and from 10% to 35% in the *RuvBL2* RNAi line (Fig. 5A). Moreover, in both induced cell lines, we observed a considerable percentage of cells with non-canonical phenotypes such as >2K2N and 1K2N (Fig. 5). At 36 h post-induction, the populations were composed of 20% and 15% of >2K2N in the *RuvBL1* and *RuvBL2* RNAi lines, respectively, and 18% and 15% of 1K2N cells. The dramatic accumulation of cells with two nuclei and multiple kinetoplasts suggested that down-regulation of either RuvB-like helicase interferes with cytokinesis, which was confirmed by fluorescence microscopy (Fig. 5B).

**Figure 5.**
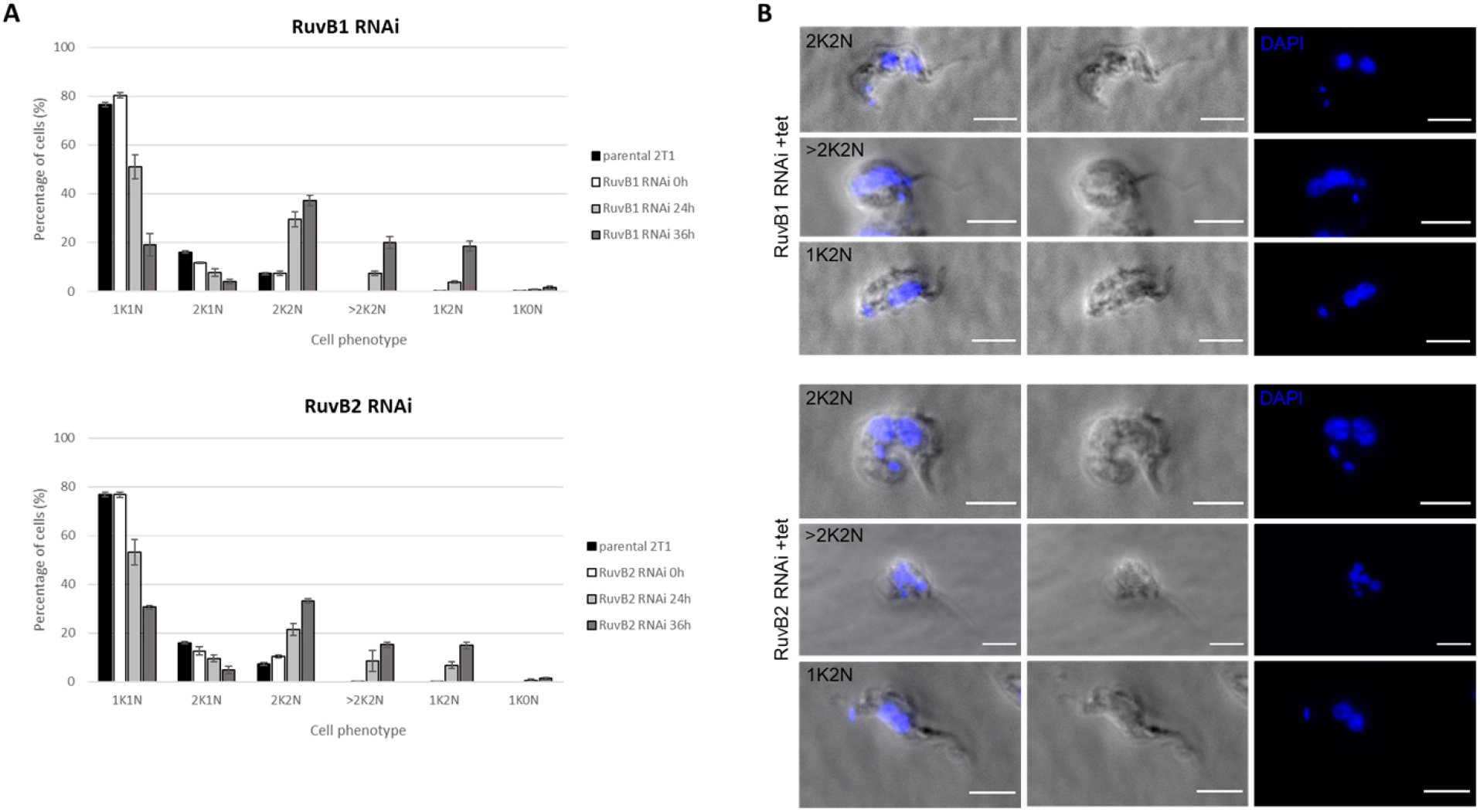
RuvBL1 and RuvBL2 are essential for cytokinesis. (A) Cell phenotype counting of *T. b. brucei* parental (2T1), uninduced (0 h), and induced clones at 24 h and 36 h post tetracycline induction (n = 3, with at least 150 cells/count). (B) DAPI staining of induced clones at 36 h post tetracycline induction; scale bar is 5 μm. Upper panel, transfectant clone D3 – *RuvBL1* RNAi; lower panel, transfectant 6B – *RuvBL2* RNAi.

## Discussion

In bacteria the RuvB-like DNA helicases are part of the resolvasome, a multiprotein complex that mediates branch migration during homologous recombination of DNA and resolution of the four-way Holliday junction (Wyatt, 2014). As such, RuvB-like helicase has a DNA-binding domain and an ATPase domain. It forms a hexameric ring around the DNA and functions as an ATP-driven motor that promotes heteroduplex formation (Wyatt, 2014). RuvB-like DNA helicases have been well studied in bacteria (Shinagawa, 1991; Iwasaki, 1992; Yamada, 2001). In eukaryotes, the proteins RuvB-like 1 and RuvB-like 2 are also called pontin and reptin (Jha, 2009), respectively, and they have acquired additional functions including DNA repair and chromatin remodeling (Tammana, 2017; Dauden, 2021). A chromatin remodelling complex from *T. brucei*, which contains TbRuvBL1 and TbRuvBL2, has been described recently (Vellmer, 2022).There is to our knowledge only one experimental study from trypanosomatids, where LmRUVBL1 and LmRUVBL2 from *Leishmania major* were recombinantly (co-)expressed. The purified proteins formed an elongated heterodimer with ATPase activity *in vitro*, but they did not form a ring (Abrahao, 2021).

Here we provide forward as well as reverse genetic evidence that RuvBL1 DNA helicase from *T. brucei* is involved in suramin resistance and a likely target of suramin: the point mutation I312V in RuvBL1 emerged heterozygously in *T. b. rhodesiense* upon suramin pressure and turned homozygous during further selection. Consistently, introduction of the mutation I312V *in situ* to RuvBL1 of *T. b. brucei* caused a 1.8-fold (heterozygous) to 2.4-fold (homozygous) increase in the IC_50_ to suramin. These resistance phenotypes were not very strong, but given the polypharmacology of suramin any mutation in just one of its targets is expected to have only a moderate effect. However, the fact that the mutation of RuvBL1 significantly lowered the susceptibility to suramin, strongly suggests that the DNA helicase is a major target of suramin in *T. brucei*. Further, this hypothesis is in agreement with the published findings that (i) suramin inhibits DNA helicases from Dengue virus (Basavannacharya, 2014), hepatitis C virus (Mukherjee, 2012), and SARS-CoV2 virus (Zeng, 2021); and (ii) exposure of bloodstream-form *T. brucei* to suramin caused a cytokinesis phenotype (Thomas, 2018) similar to the one observed upon RNAi-mediated down-regulation of RuvBL1. Functional assays to corroborate a direct interaction between *T. brucei* RuvBL1 helicase and suramin will be challenging, requiring the presence of RuvBL2 and possibly further proteins of the complex, or even a DNA substrate.

Mutation of RuvBL1 isoleucin-312 was also detected in a suramin-resistant field isolate of *T. evansi*, in that case to leucin. *T. evansi* is a subspecies of *T. brucei* (Lai, 2008) that is mechanically transmitted by biting flies and is no longer dependent on the tsetse fly host. Thus the parasite has spread across the tropical regions globally (Desquesnes, 2013), and cases were reported even from southern Europe (Desquesnes, 2008; Tamarit, 2010). The resistant *T. evansi* strain Westry 2 was isolated from an infected camel in the Republic of the Sudan. It had an IC_50_ to suramin of over 40 μg/ml (28 μM) as determined in a short-term 3H-hypoxanthine incorporation assay (El Rayah, 1999). This isolate had not been adapted to *in vitro* culture, preventing its further characterization. Given the polypharmacologic nature of suramin, drug resistance is more likely to arise by transport phenotypes (increased export or decreased import) than by target site mutation. Nevertheless, the mutation of the conserved isoleucin-312 in a field isolate indicates RuvBL1 relevance for suramin resistance in the field, warranting further investigation of RuvBL1 in *T. evansi*.

Sequence analysis and the TbRubBL1 homology model places I312 in close vicinity to the active site and motifs with crucial residues engaging in ATP binding and hydrolysis, suggesting the possibility of suramin inhibition via binding to the respective region. Further, I312 is located near the pore of the hexameric assembly upstream the sensor 1 motif (Fig. 3B, Fig. 3C), a region likely interacting with DNA substrate. Here, suramin binding could interfere with substrate interaction or even trigger substrate sensing-dependent, perpetual ATPase activity. However, I to L and I to V mutations are considered conservative, retaining a hydrophobic, rather less bulky side chain, rendering a direct effect on suramin binding less likely. It is also possible that the I312 mutation leads to a structural rearrangement that in turn effects interaction with suramin. Altogether, the precise molecular mechanism of suramin interaction with TbRuvBL1 remains to be investigated.

Finally, our findings render RuvBL1 an attractive drug target. It was recently proposed as a novel target for antimalarials (Khurana, 2021). The facts that (i) RNAi-mediated silencing of RuvBL1 expression in bloodstream-form *T. brucei* was rapidly lethal, and (ii) the RuvBL1 orthologues from kinetoplastids form a distinct phylogenetic clade, qualify RuvBL1 as a target in drug development for trypanosomatid diseases.

## Methods

### Trypanosoma spp. strains

*Trypanosoma b. rhodesiense* STIB900_c1 was a fresh clone of STIB900 (Wiedemar et al., 2017), which derived from STIB704 isolated in 1982 from a male patient in St. Francis Hospital, Ifakara, Tanzania. *Trypanosoma b. brucei* 2T1 is a derivative of Lister427 and is widely used for reverse genetics (Alsford, 2005). *Trypanosoma evansi* Westry2 and STIB779 were isolated from camels in Western Sudan in 1995 and Kenya in 1979, respectively (El Rayah, 1999).

### Cell culture

Parasites were cultivated at 37°C, 5% CO_2_ in Iscove’s Modified Dulbecco’s Medium supplemented according to Hirumi (Hirumi, 1989) and with 15% heat-inactivated horse serum for *T. b. rhodesiense* or 10% heat-inactivated fetal calf serum for *T. b. brucei* 2T1. The selection for suramin resistance was carried out under constant drug pressure with gradually increasing concentrations from 1.1 μM to 18.9 μM suramin over the course of one year (Fig. 1).

### Drug testing

Drug efficacy was determined *in vitro* with the Alamar Blue assay (Räz, 1997). Serial drug dilutions were prepared in a 96-well plate, and parasites were added to a concentration of 2 × 10^4^ cells/ml for STIB900 and 10^4^ cells/ml for 2T1. After 68 h of incubation, resazurin was added at a concentration of 11.4 μg/ml, and the plates were incubated for 2 to 4 more hours. The fluorescence was quantified with a SpectraMax reader (Molecular Devices) and SoftMax Pro 5.4.5 Software. GraphPad Prism 6.0 was used to fit dose-response curves (non-linear regression model, variable slope, four parameters, lowest value set to zero) and to calculate the IC_50_ values.

### Isolation of nucleic acids, polymerase chain reaction

Genomic DNA for PCR and Sanger sequencing was isolated from approximately 10^7^ bloodstream-form *T. brucei*, grown *in vitro* and washed once with PBS, using the DNeasy Blood and Tissue Kit (Qiagen) according to the manufacturer’s protocol. The DNA was eluted in 200 μl milliQ water. PCR was carried out with the KAPA HiFi PCR Kit (KAPA Biosystems) in a total volume of 25 μl. Primers for amplification and sequencing of Tb927.4.1270 a934g were Tb927.4.1270-F-01 and Tb927.4.1270-R-01-BamHI; all primers are listed in Supplementary Table S1. Genomic DNA for genome sequencing of *T. evansi* was isolated from cells propagated in mice (female NMRI, 22 to 25 g). The trypanosomes were collected from euthanized mice by heart puncture, separated from the blood with a diethylaminoethyl cellulose column prepared from Whatman DEAE cellulose (DE52 pre-swollen, 3W4057-200) (Lanham, 1970), and the genomic DNA was isolated by chloroform/phenol extraction.

RNA was isolated from approximately 10^7^ cells grown to a density of 10^6^ cells/ml using the RNAeasy Mini Kit (Qiagen), including an on-column DNase I treatment (Qiagen). The RNA was isolated from each cell line in triplicates from independent cultures. For RT-PCR, *T. b. brucei* 2T1 lines (RuvBL1 RNAi, RuvBL2 RNAi) were grown for 24 h in the absence or presence (1 ug/ml) of tetracycline. For RNA-Seq, the four *T. b. rhodesiense* STIB900 lines (c1, c1_sur1, c1_sur1_3500, c1_sur1_18900) were grown in parallel without suramin.

### RNA-Seq and analysis

Sequencing libraries for RNA-Seq were prepared separately for each replicate using the TruSeq Stranded mRNA Library preparation kit (Illumina). 126-nucleotide single-end sequencing was carried out on an Illumina HiSeq 2500 machine. The quality of the reads was determined with fastqc (Andrews, 2010). Since the quality was high, the reads were mapped untrimmed to the *T. b. brucei* TREU927 reference sequence (TryTrypDB-38), supplemented with the Lister427 bloodstream expression sites (Hertz-Fowler, 2008) plus *VSG^900^* (GenBank MF093646) and *VSG^Sur^* (GenBank MF093647), with the Burrows-Wheeler Aligner (Li, 2009a) using the default settings. The sam-files were converted to bam-files using samtools (Li, 2009b) and sorted and indexed with Picard (Institute, 2019); read duplicates were marked and read group identifiers added with Picard as well. Deviations from the reference sequence (variants) were called with the GATK version 4.0.7.0 haplotypecaller (McKenna, 2010) in gvcf-mode. The individual g.vcf-files were combined into one using GATK combinegvcfs and genotyped using GATK genotypegvcfs in order to obtain a VCF-file with all the identified variants. Variant annotation was calculated using snpEff version 4.3T (Cingolani, 2012) based on the TREU927 reference annotation (TryTrypDB-38) complemented with the genes of the Lister427 bloodstream expression sites, *VSG^900^*, and *VSG^Sur^*. Variants were excluded that laid within highly diverse regions with a low quality of alignment, were wrongly called due to small differences in allele frequencies, or were within VSG genes or pseudogenes. Sequencing data were deposited in the European Nucleotide Archive (accession PRJEB51200).

### Whole genome sequencing

DNA Sequencing libraries were prepared using Illumina’s KAPA Hyper Prep Kit, and paired-end 126 nts sequencing was carried out on an Illumina HiSeq 2500 machine. Raw sequencing reads were mapped to the *T. evansi* reference genome STIB805 (TriTrypDB-46) using the Burrows-Wheeler Aligner (Li, 2009a). The sam-files were converted to bam-files using samtools (Li, 2009b), then sorted and indexed using Picard (Institute, 2019). The aligned sequencing reads were visualized with the integrative genomics viewer (Robinson, 2011; Thorvaldsdottir, 2013) to inspect the candidate genes that had been detected in the *T. b. rhodesiense* STIB900 derivatives. Sequencing data were deposited in the European Nucleotide Archive (accession PRJEB51200).

### Reverse genetics with T. brucei

The construct for introduction of the mutation a934g into *RuvBL1* was synthesized (GenScript, Netherlands). It contained the last 704 nucleotides of the Tb927.4.1270 coding sequence including the mutation, the 3’UTR, an βα tubulin mRNA processing site, a blasticidine resistance gene, an αβ tubulin mRNA processing site, and the 215 nucleotides downstream of Tb927.4.1270. For the generation of homozygous mutants, the blasticidine resistance gene was exchanged with a neomycin resistance gene. The construct was amplified by PCR (primers Tb927.4.1270-F-01 and Tb927.4.1270-R-02; Table S1) and purified using the NucleoSpin® Gel and PCR Clean-up kit (Macherey-Nagel). 3 to 4×10^7^ cells were transfected with 3 to 8 μg of DNA in 100 μl bloodstream form transfection buffer (Schumann Burkard, 2011) using program Z-001 on the Amaxa Nucleofector (Lonza). Limiting dilutions of the parasites were prepared in 48-well plates. After one day, selection antibiotics were added: blasticidine (Invivogen) at 5 μg/ml for *T*. *b. rhodesiense* and 10 μg/ml for 2T1 cells; G418 (Invivogen) initially at 3 μg/ml, reduced to 1 μg/ml after picking of clones 4 to 5 days after transfection.

For overexpression of *RuvBL1* and *RuvBL2*, a DNA fragment containing both coding sequences with an interspaced αβ-tubulin mRNA processing site and flanked with HindIII and BamHI sites was synthesised (GenScript, Netherlands) with re-coded internal HindIII sites to allow cloning. For simultaneous overexpression of *RuvBL1* and *RuvBL2*, the fragment was directly cloned into a pRPa plasmid. For overexpression of *RuvBL1* alone, the re-coded *RuvBL1* sequence was amplified by PCR (primers Tb927.4.1270-F-02-HindIII and Tb927.4.1270-R-03-BamHI), digested, purified, and cloned into a pRPa plasmid. All constructs were checked by Sanger sequencing. 4×10^7^ 2T1 cells were transfected with approximately 5 μg of SacI digested and purified plasmid. The parasites were selected with 5 μg/mL hygromycin B Gold (Invivogen). Overexpression was induced with 1 μg/ml tetracycline (Sigma-Aldrich).

For inducible knock-down of *RuvBL1* or *RuvBL2* expression, the stem-loop pRP-iSL MSC1/2 plasmid was used to generate two RNAi constructs. Optimal targeting sequences (i.e. nts 483-1076 for *RuvBL1* and 236-749 for *RuvBL2*) as identified by the RNAit tool (Redmond, 2003) were amplified by PCR with overhang primers (Tb927.4.1270-F-03 and Tb927.4.1270-R-05; Tb927.4.2000-F-02 and Tb927.4.2000-R-02) using the re-coded genes as template. After PCR clean-up and digestion, the sense fragment was cloned into the pRP-iSL plasmid, followed by insertion of the antisense fragment. All constructs were checked by Sanger sequencing. The plasmids were digested with AscI, and purified prior to transfection, as described above. Transfectants were selected with 5 μg/mL hygromycin B Gold (Invivogen). RNAi was induced with 1 μg/mL tetracycline (Sigma).

### cDNA synthesis and quantitative PCR

Complementary DNA was synthesized with the SuperScript™ III Reverse Transcriptase kit (Invitrogen) using an oligo(dT)_15_ primer (Promega). Quantitative PCR (primers Tb927.4.1270-F-01 and Tb927.4.1270-R-04; Tb927.4.2000-F-01 and Tb927.4.2000-R-01) was carried out in duplicates using Fast SYBR® Green Master Mix (Applied Biosystems) in a StepOnePlus real-time PCR System (Applied Biosystems). Data were analyzed with StepOne Software v2.3. Telomerase reverse transcriptase (*TERT*) was used as a housekeeping gene for normalization with the ΔCt method.

### DAPI staining and phenotype scoring

*T. b. brucei* 2T1 *RuvBL1* and *RuvBL2* RNAi parasites, uninduced or induced for 24 h and 36 h with 1 μg/mL tetracycline, were spun down for 5 min at 800 g, washed once in 1x Voorheis-modified PBS (vPBS: 137 mM NaCl, 3 mM KCl, 16 mM Na_2_HPO_4_, 3 mM KH_2_PO_4_, 46 mM sucrose, 10 mM glucose, pH 7.6), and resuspended in 1% PFA (EMS). The cells were incubated for 2 min on ice, centrifuged for 5 min at 2000 g, and the pellet was resuspended in 1x PBS. Cells were loaded on microscopy slides (Hecht Assistent), allowed to adhere for 10 min, then permeabilized in ice-cold MeOH for 30 min. Nuclei and kDNAs were stained for 2 min with 10 μg/mL DAPI (Sigma), washed, mounted with VECTASHIELD® antifade mounting medium, and covered with coverslips (Duran). Cells were imaged with a Leica DM5000B microscope equipped with a Leica K5 Camera using the LAS v4.9 software. Images were processed with ImageJ Fiji. At least 150 cells per condition, in triplicates, were counted.

### Homology modeling

Homology models of TbRuvBL1 (Tb927.4.1270) and TbRuvBL2 (Tb927.4.2000) were built with Phyre2 (Kelley, 2015) using *Saccharomyces cerevisiae* RuvBL1 (pdb ID 6GEN) (Willhoft, 2018) as template, yielding confidence scores of 100%, respectively. Structural alignments were performed, and structural models visualized in PyMOL (Schrödinger, 2020).

### Phylogeny

Amino acid sequences were acquired from different taxonomic groups by Blastp searches (Altschul, 1990) using *T. brucei* RuvBL1 as the query. Multiple sequence alignment (Edgar, 2004), construction of a Neighbour-Joining tree (Saitou, 1987), and boostrapping (Felsenstein, 1985) were done with MegaX (Kumar, 2018). The alignment was trimmed at the N- and C-terminus for overhanging sequences. Distances were computed using the JTT matrix (Jones, 1992). The alignment in Figure S2 was made with Clustal Omega (Sievers, 2011) and edited with Jalview version 2.1.1.4 (Waterhouse, 2009).

## Acknowledgments

We wish to thank Sam Alsford and Matthew Gould for kindly providing the pRP plasmids, Kirsten Gillingwater and Christina Kunz Renggli for infecting mice with *T. evansi* and harvesting the trypanosomes, and the Genomics Facility Basel for high-throughput sequencing. Genomics analyses were performed at sciCORE scientific computing center of the University of Basel (http://scicore.unibas.ch/). This study was financially supported by the Swiss National Science Foundation (grant 310030_156264 to PM) and the Czech Ministry of Education (grant OPVVV/0000759 to MZ).

---

**Table S1.**
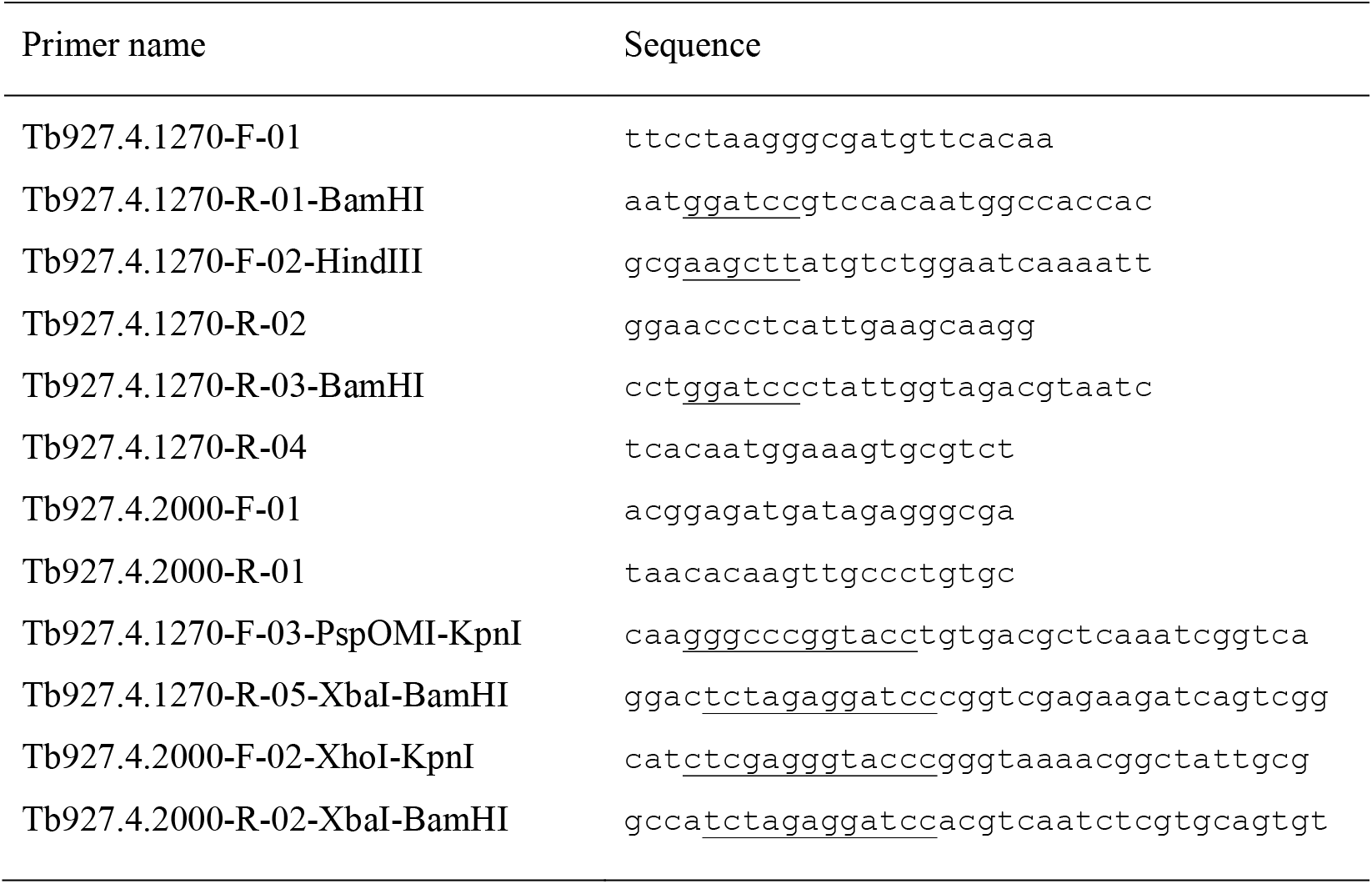
Primers used for PCR. All sequences are 5′ to 3′. Restriction sites are underlined.

**Supplementary Figure S1.**
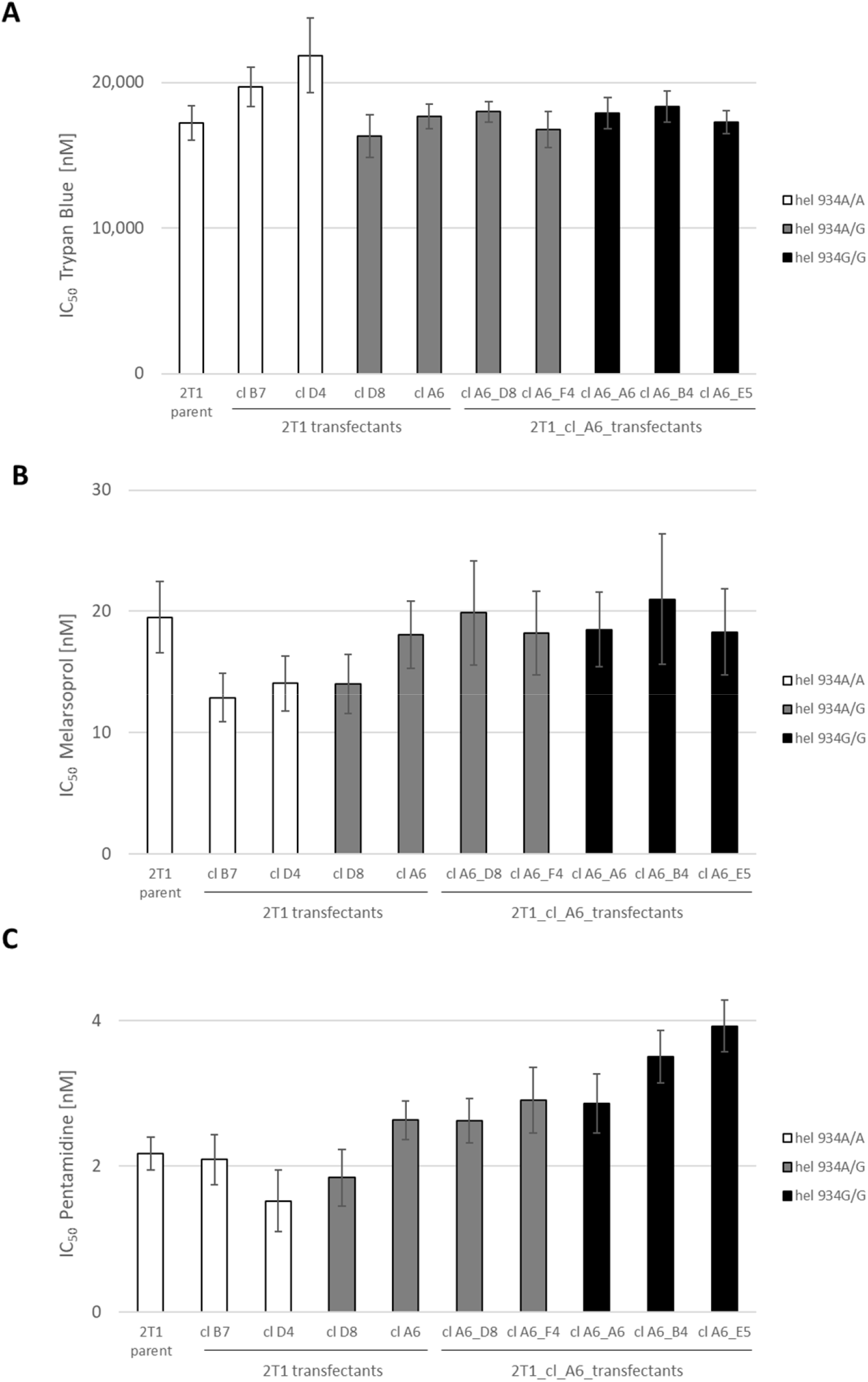
Drug sensitivity of *T. b. brucei* 2T1 after *in situ* introduction of the mutated *RuvBL1* I312V coding sequence. 50% inhibitory concentrations (IC_50_) of trypan blue (A), melarsoprol (B), and pentamidine (C) to the recovered transfectants after the first round of transfection (4 clones: wildtype in white, heterozygous mutants in grey) and the second round (5 clones: heterozygous mutants in grey, homozygous ones in black).

**Supplementary Figure S2.**
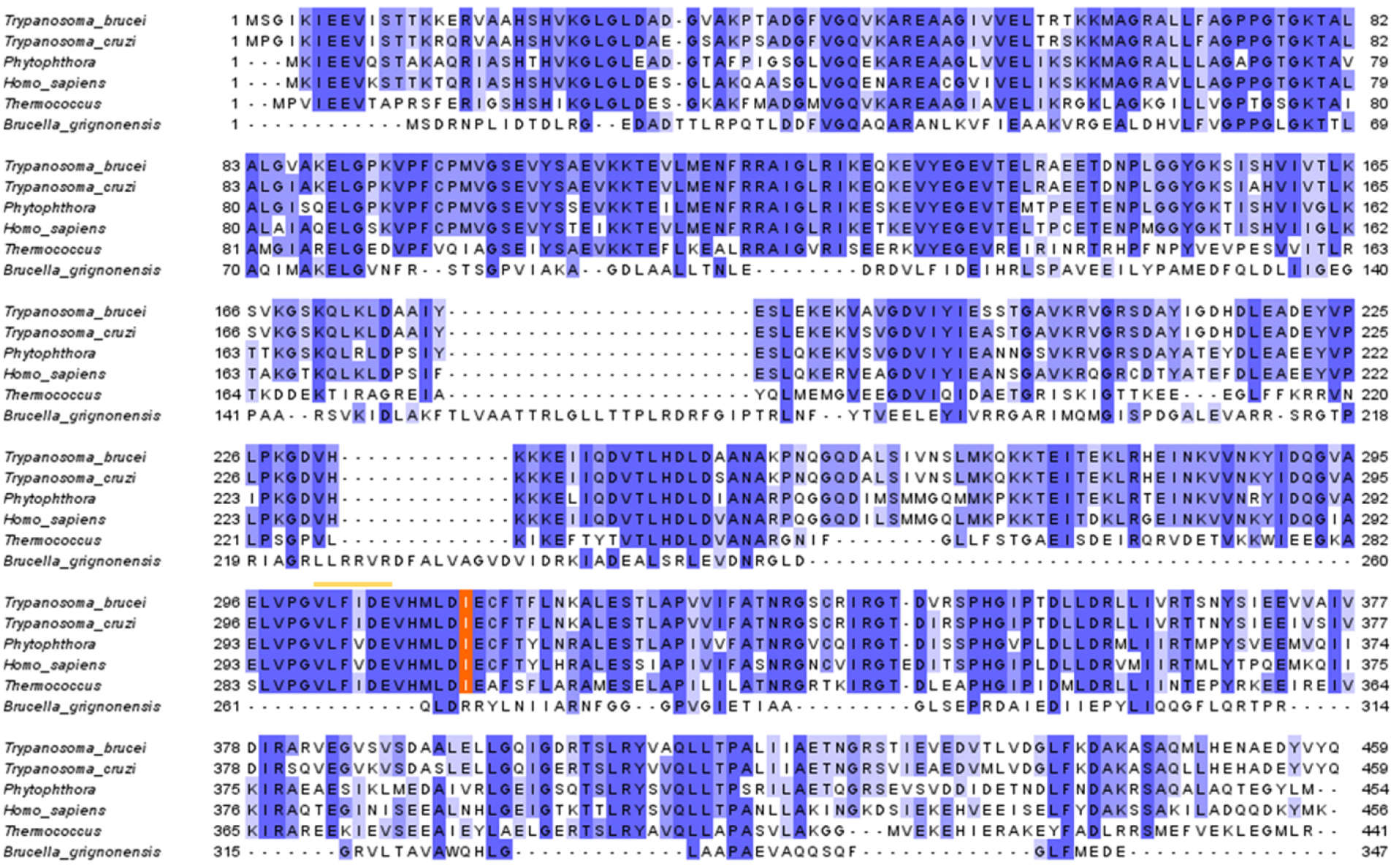
Multiple alignment of RuvB-like helicases. RuvB1 helicase is conserved among kinetoplastids (*T. b. brucei* and *T. cruzi*) and beyond (i.e. oomycetes: *Phytophthora rubi*, animals: *Homo sapiens*, and archaea: *Thermococcus*), but not bacteria (*Brucella grigionensis*). The orange box marks the isoleucin in position 312. The color-code blue-to-white indicates the percentage of identity of the single amino acids between the six protein sequences. The yellow line marks the conserved Walker B motif (hhhhDE) (Hanson, 2005; Matias, 2006). The multiple sequence alignment was edited with Jalview (Waterhouse, 2009).

**Supplementary Figure S3.**
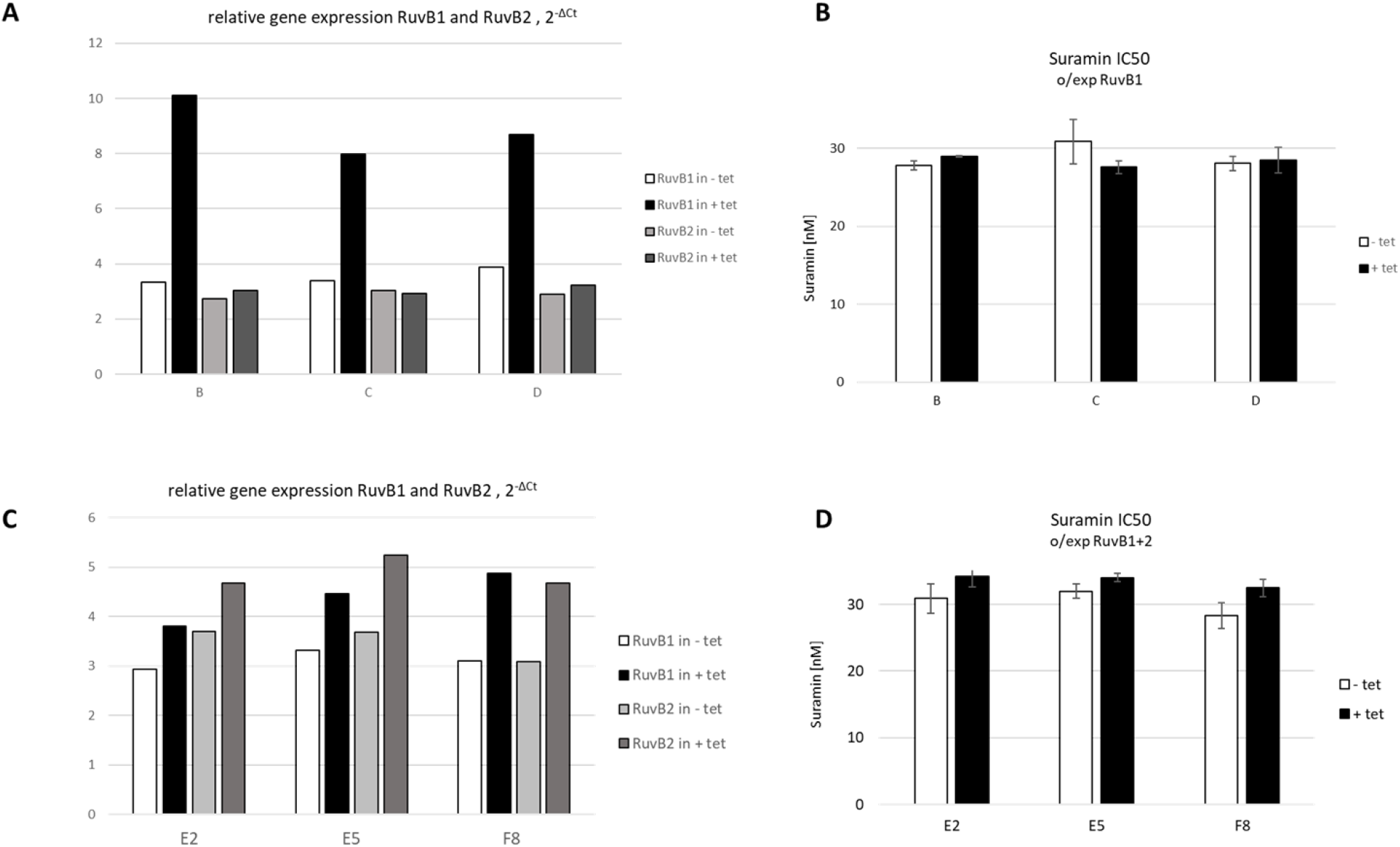
Overexpression of *RuvBL1* in bloodstream-form *T. brucei*. (A) Gene expression levels as determined with qPCR (2 technical replicates) for three *RuvBL1* over-expressing clones (B, C, D). (B) Suramin sensitivity of induced and non-induced clones (n = 3) over-expressing RuvBL1. (C) Gene expression as determined with qPCR (2 technical replicates) for three RuvBL1+2 over-expressing clones (E2, E5, F8). (D) Suramin sensitivity of induced and non-induced clones (n = 3) over-expressing RuvBL1+2.

